# Unraveling new characteristics of γδ T cells using scRNA-seq in TCR KO chicken

**DOI:** 10.1101/2025.02.26.640441

**Authors:** Theresa von Heyl, Milena Brunner, Antonia Schmauser, Christine Wurmser, Simon Früh, Benjamin Schusser

## Abstract

The characterization of T cells in chickens has proven to be challenging, primarily due to the lack of specific markers to differentiate classical Th1, Th2, and Th17 subsets. Furthermore, chickens possess a notably high proportion of γδ T cells, making them a unique model for investigating the poorly understood role of these cells not only in chickens but also in mammals. To gain deeper insights into the functions and characteristics of the αβ and γδ T cell subsets in chickens, whole transcriptome analysis (WTA) on CD3^+^ single cells isolated from wild type (WT), TCR Cβ knockout (KO) and TCR Cγ KO chickens at embryonic day (ED) 18, day (d) 3, and d14 post-hatch was conducted. The results indicate that γδ T cells exhibit cytolytic activity in both TCR Cβ KO and WT chickens. A distinct cluster of γδ T cells expressing genes associated with the interferon pathway was revealed in TCR Cβ KO chickens, which may contribute to the severe phenotype in these chickens, which is characterized by severe inflammation of spleen, gut and stomach. Additionally, novel gene markers were identified that precisely define the αβ and γδ T cell subsets. These findings provide new insights into chicken T cell biology and contribute to a better understanding of the mechanisms underlying the severe phenotype observed in TCR Cβ KO chickens.

## 1 Background

Chickens are an indispensable livestock species, as not only the chicken meat but also the eggs are a principal protein source worldwide. Compared to other livestock species, chicken meat is a healthier and more resource-efficient source of protein (Farrell 2011, Salter 2018). Livestock diseases often pose significant threats, leading to great economic losses. Moreover, diseases like avian influenza pose a considerable zoonotic risk to humans (Bodewes, Bestebroer et al. 2015, Rahman, Sobur et al. 2020, Abdelwhab and Mettenleiter 2023). Therefore, a detailed understanding of the chicken immune system is essential to ensure the health of chickens and humans.

T cells are key players in the adaptive immunity of chickens (Cooper and Alder 2006). They are categorized into αβ and γδ T cells based on their T cell receptor chains (Chen, Sowder et al. 1989). The role of αβ T cells is defined through co-expression of CD4 and CD8 (Arstila, Vainio et al. 1994, Luhtala 1998). The CD4^+^ αβ T cells are activated through antigen peptides presented on MHC class II molecules, leading to production of cytokines, targeted immune responses and the stimulation of B cells to secrete immunoglobulins (Arstila, Vainio et al. 1994). The CD8^+^ T cells can directly lyse pathogens by secreting cytotoxic cytokines upon activation through peptides on MHC class I molecules (Luhtala 1998). The balance between protective inflammation and tissue damage is stabilized by regulatory T cells (T_reg)_, described by their transcription factor *FoxP3* (Burkhardt, Elleder et al. 2022). Compared to mammals, there is no clear classification into Th1, Th2 and Th17 cells of αβ T cells in chickens, as the classification is based on the secretion of various interleukins, some of which have not yet been described (Mousset, Hobo et al. 2019).

The other major T cell subset, γδ T cells, remains poorly understood in both chickens and mammals. Chickens present a compelling model for studying the role of γδ T cells in greater detail, as they are a γδ-high species, with comprising up to 50% γδ T cells of circulating lymphocytes throughout their lifespan (Sowder, Chen et al. 1988). In contrast, mice and humans typically harbor only 1–10% γδ T cells in the peripheral blood(Sowder, Chen et al. 1988, Holderness, Hedges et al. 2013). To date, chicken γδ T cells are known to exhibit cytotoxic activity when stimulated *in vitro* (Fenzl, Gobel et al. 2017). They are abundantly present in peripheral epithelial tissues (Hayday, Roberts et al. 2000), playing a crucial role in the immune response to infections such as Salmonella (Berndt and Methner 2001). γδ T cell subsets have been previously characterized by Berndt et al. into three groups: CD8^high+^, CD8^dim+^, and CD8^neg-^. These γδ T cell subsets respond differently to infections like salmonellosis (Berndt, Pieper et al. 2006).

To characterize T cell subsets in more detail, we generated knockout chickens that lack αβ or γδ T cells, respectively. The phenotypical analysis of the TCR Cγ KO chicken showed a comparable phenotype to wild type (WT) chickens (von Heyl, Klinger et al. 2023). Additionally, infection of TCR Cγ KO chickens with MDV showed increased disease incidence and tumor formation compared to their WT siblings, leading to the conclusion that γδ T cells play an important role in the pathogenesis and tumor formation (Sabsabi, Kheimar et al. 2024). Strikingly, TCR Cβ KO chickens exhibit a severe phenotype two weeks after hatch, with granuloma formation at the comb, beak, and legs. In addition, inflammation of the stomach and spleen, as well as an impaired development of the lymphatic organs was observed. The cause of this phenotype and a potential contribution of the remaining γδ T cells remain elusive (von Heyl, Klinger et al. 2023).

Using scRNA-seq analysis, Maxwell et al. identified leukocyte populations such as B cells, eosinophils, basophils, monocytes, red blood cells, and thrombocytes in adult 24 week old laying hens. The description of T cell subtypes proved to be challenging in chickens, but it led to new insights regarding cytotoxic characteristics of particular T cells (Maxwell, Soderlund et al. 2024).

Here, we used scRNA-seq to perform whole transcriptome analysis (WTA) of CD3^+^ T cells sorted from the thymus and peripheral blood mononuclear cells (PBMCs) of WT, TCR Cβ KO and TCR Cγ KO animals on embryonal day 18 (ED18), day 3 (d3) and day 14 (d14) post hatch. Our data revealed two different cytotoxic γδ T cell clusters in the TCR Cβ KO at 14 days post-hatch, which were absent in TCR Cγ KO and WT chickens as well as potential gene markers to identify γδ and αβ T cell clusters.

## 2 Methods

### 2.1 Animals

Two genetically modified chicken lines, TCR Cβ KO and TCR Cγ KO, derived from the White Leghorn genetic background (Lohmann Selected Leghorn [LSL]; Lohmann-Tierzucht GmbH, Cuxhaven, Germany), were used in this study. The generation and phenotypic characterization of these lines have been described previously (von Heyl, Klinger et al. 2023). Animal experiments were approved by the government of Upper Bavaria, Germany (ROB-55.2-2532.Vet_02-17-101 & 55.2-1-54-2532-104-2015). Experiments were performed according to the German Welfare Act and European Union Normative for Care and Use of Experimental Animals. All animals received a commercial standard diet and water *ad libitum*.

### 2.2 Organ / Tissue collection

For the sampling of the thymus of ED18 chicks, TCR Cγ KO and TCR Cβ KO eggs were incubated for 18 days in egg incubators (HEKA-Brutgeräte, Rietberg, Germany) at standard conditions (37,8°C. 55% relative humidity, six rotations/day). Infertile eggs or dead embryos were removed by candling the eggs on day 7 of incubation. *In ovo* genotyping was performed on ED10. On ED18, eggs were opened, and embryos were decapitated and the thymus was collected from the embryos. For the other time points, TCR Cβ KO, TCR Cγ KO, and their WT sibling were hatched. Blood was drawn for genotyping on d1 post hatch. On d3 and d14 post hatch, chickens were euthanized by terminal bleeding, and the thymus was removed. From all genotypes (WT, TCR Cβ KO and TCR Cγ KO) n=3 animals irrespective of sex were used at each time point (ED18, d3, d14).

### 2.3 Genotyping assays

For *in ovo* genotyping, allantois samples were collected on ED10. Briefly, a candling light was used to determine fertility of the egg, embryo viability, and to mark the areas for creating a hole with a lancet where no blood vessels were visible. The allantoic fluid was collected using an insulin syringe with a 30G cannula (BD Micro Fine^TM^, Heidelberg, Germany). Allantoic fluid was directly used for genotyping without a gDNA isolation step.

For the hatched chicks, a drop of blood was collected one day after hatch from the *vena jugularis* using a syringe with a 27G cannula (B. Braun, Meisungen, Germany). gDNA was isolated from blood, by adding 1µL blood into 200µL STM buffer and centrifuging for 5min at 100g. After removing the supernatant, 400µL TEN buffer containing 100µg/mL Pronase E was added, and the mix was incubated for 1h at 37°C (250rpm). The Pronase E was subsequently inactivated at 65°C for 20min (Bailes, Devers et al. 2007).

Chickens were genotyped as described before (von Heyl et al.). Briefly, primers were used to detect TCR Cβ KO chickens: Forward: 5’ GGTTCGAAATGACCGACCAAGC 3’; Reverse: 5’ GGCTTGCACACTCAGCTCTATAG 3’. A second primer pair was used to detect the TCR Cβ WT allele: Forward: 5’ GGTTCGAAATGACCGACCAAGC 3’; Reverse: 5’ CACACCATTCACCTTCCAGAC 3’. FIREPol Multiplex DNA Polymerase Mastermix (Solis Biodyne, Tartu, Estonia) was used according to manufactures instructions with an annealing temperature (Tm) of 58°C. For the TCR Cγ KO chickens’ primers were used to detect the TCR Cγ KO: Forward: 5’ GCCATTCCTATTCCCATCCTAAGT 3’; Reverse: 5’ GGTTCGAAATGACCGACCAAGC 3’. A second primer pair was used to detect the WT constant region of the TCR γ chain: Forward: 5’ GAGCTCCACGCCATGAAACCATAG 3’; Reverse: 5’ GTTGTCACTGTCACTGGCTG 3’. FIREPol Multiplex DNA Polymerase Mastermix (Solis Biodyne, Tartu, Estonia) was used according to manufactures instructions with an annealing Tm of 60°C.

### 2.4 Cell sorting

The removed thymus was strained through a 100µm cell strainer into a falcon holding 5mL ice-cold PBS. Lymphocytes were isolated from blood and the strained thymus single-cell solution using histopaque density gradient centrifugation (Sigma, Taufkirchen, Germany). Cells were washed with 1% BSA in PBS and stained with CD3-PE (clone CT-3; 0,5µg/mL) (Biozol, Eching, Germany) for 20 min in the dark on ice. Subsequently, cells were again washed, and cell sorting was performed using a CytoFLEX SRT Cell sorter (BD, Heidelberg, Germany). Data were analyzed with FlowJo 10.8.1 software (FlowJo, Ashland, USA).

### 2.5 Multiplexing

In order to multiplex all samples from each timepoint the BD Flex Single-Cell Multiplexing Kit A, B, and C (BD, Heidelberg, Germany) were used according to the manufacturer’s protocol. Briefly, cells were resuspended in 180µL PBS+ 1%FBS. 20µL of each sample tag was added and incubated. Samples were washed with PBS+ 1%FBS and pooled together.

### 2.6 Single-cell capture and cDNA Synthesis

The BD Rhapsody**™** cDNA Kit and BD Rhapsody Enhanced Cartridge Reagent Kit (BD, Heidelberg, Germany) were used according to the manufacturer’s protocol. The 8-Lane Cartridge was loaded with 70.000 cells.

### 2.7 mRNA Whole Transcriptome Analysis and Sample Tag

The BD Rhapsody™ WTA Reagent Kit (BD, Heidelberg, Germany) was used according to the manufacturers protocol (23-24119(02)). Briefly, random priming and extension (RPE) reactions of the Exonuclease I-treated beads containing the whole cellular transcriptome and cDNA was performed. After this step two parallel workflows were performed, one for the WTA libraries, using the RPE product from the supernatant and the other one for the Sample Tag libraries from the post-RPE beads. The RPE supernatant underwent WTA library preparation following the manufacturers’ instructions from the BD Rhapsody™ System mRNA Whole Transcriptome Analysis (WTA) Library Preparation Protocol that included the steps of RPE cleanup, performing RPE PCR, RPE PCR cleanup, performing WTA Index PCR and cleanup. The Sample Tag products were amplified and purified through PCR. A Sample Tag Index PCR was performed to add full-length Illumina Sequencing adapters and indices.

WTA libraries and Sample Tag libraries were pooled and sequenced on a NovaSeqX 10B Sequencer by Novogene (Cambridge, UK). Libraries were sequenced as paired-end 2x150bp reads with an estimated depth of 1,250 M reads per cartridge lane.

### 2.8 Single-cell data analysis

#### 2.8.1 Preprocessing of scRNA-seq raw data

Fastq files were processed using the BD Rhapsody™ WTA Analysis Pipeline, v1.9.1 in the Seven Bridges Genomics cloud platform. WTA reads were aligned to the *Gallus gallus* (chicken) GRCg7b reference genome (Accession Number GCF_016699485.2) and demultiplexed. The pipeline generates molecular counts per cell, reads per cell, metrics and an alignment file.

#### 2.8.2 Integration, Clustering and Annotation

Bioinformatic analyses were conducted in RStudio (Version 2024.04.2+764) using Seurat (Version 5.1.0). Cluster analyses were performed following the Seurat integration workflow (Version 5.2.0; https://satijalab.org/seurat/articles/integration_introduction). Differentially expressed genes (DEGs) were identified from the integrated clusters according to the Seurat pipeline (https://satijalab.org/seurat/articles/pbmc3k_tutorial.html). Functional classification of gene markers was carried out using PANTHER (Version 19.0) for gene ontology (GO) analysis. Cluster annotation was performed manually based on the inspection of DEGs.

## 3 Results

### 3.1 Integrated Transcriptomic Landscapes of CD3^+^ T cells in Thymus and PBMCs

In this study, whole transcriptome analysis was conducted on sorted CD3^+^ T cells isolated from WT, TCR Cβ KO, and TCR Cγ KO animals. Thymus samples were collected from ED18, d3, and d14 chickens, while PBMC samples were obtained from d3 and d14 chickens (Figure 1A). Due to the limited blood volume in ED18 embryos, it was not feasible to sort the required 20,000 cells per embryo. Generally, the CD3^+^ population in the thymus ranged from 15% to 49,4% at all timepoints. The CD3^+^ population in blood increased from d3 to d14. At d3 the population ranged between 0,73% to 8,98% of all PBMCs with no differences between the genotypes. At d14 TCR Cβ KO chickens showed a smaller population of CD3^+^ PBMCs (mean 2,57%), compared to TCR Cγ KO (mean 8,88%) and WT (mean 10,17%) animals (Figure 1B and C and Supplement Figure 1-3). From each sample 20,000 cells were sorted, except for one TCR Cβ KO thymus sample at d14 where only 8,000 sorted cells were achieved. The quality of the sorting captured using BD Rhapsody^TM^ Scanner was summarized for each timepoint in Table 1 and 2. The Sequencing quality was overall good since over 96% of the reads passed the quality filter and over 90% of the reads were assigned to a putative cell at all three timepoints (Table 1 and 2). At all three timepoints over 90% of the reads were aligned to a putative cell. At d3 the highest mean with 14,650.78 reads per cell was found. On ED18 39,080, on d3 45,884 and on d14 80,075 viable cells were loaded. The cell multiplet rate ranged from 7,2% to 15,8% (Table 1). The single cell captures overall showed a good viability of the cells with a bead efficiency of 94,6%-98,2% (Table 1). Clusters from all timepoints, organs and genotypes were integrated for comparison, resulting in a total of 17 identified Clusters. In general, the clusters of TCR Cβ KO and TCR Cγ KO showed only few overlaps in the thymus samples (Figure 3 A-C).

**Figure 1.**
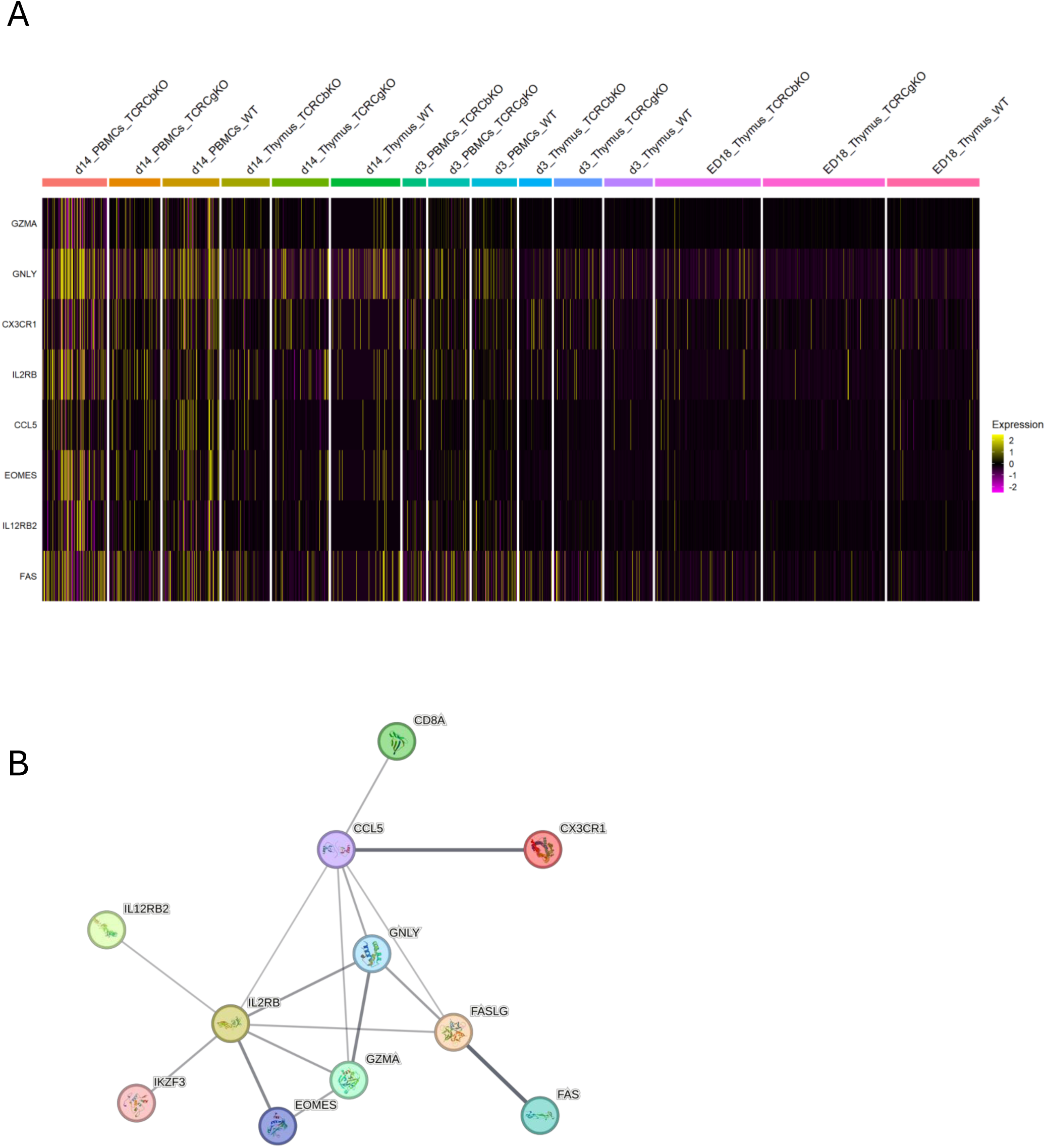
Experimental setup for the single cell analysis of chicken CD3^+^ T cells. (A) PBMC and thymocytes of three chickens from each genotype WT, TCR Cβ KO and TCR Cγ KO were used. CD3^+^T cells were isolated by fluorescence-activated cell sorting (FACS) and applied to 5’ single-cell RNA sequencing (5’ scRNA-seq) for whole transcriptome analysis (WTA) (created with BioRender). (B) Flow cytometric analysis of CD3^+^ T cells from Thymus and PBMCs of WT, TCR Cβ KO and TCR Cγ KO chickens n=3 at ED18, d3, d14 (Mean +/- SD are shown). (C) The gating strategy for isolation of CD3^+^ T cells by FACS. One representative dotplot of each timepoint (ED1, d3, d14) and each genotype (WT, TCR Cβ KO and TCR Cγ KO) is shown.

**Table 1.** Quality control metrics of BD Rhapsody^TM^ Scanner.

| Timepoint | Number of wells with viable cells at cell load | Cell multiplet rate | Bead load efficiency |
| --- | --- | --- | --- |
| ED18 | 39080 | 7,2% | 94,6% |
| d3 | 45884 | 9,8% | 97,9% |
| d14 | 80075 | 15,8% | 98,2% |

**Table 2.** Sequencing quality of cells and sample multiplexing provided by Novogene:

|  | Cells |  |  |  | Sample Multiplexing |  |
| --- | --- | --- | --- | --- | --- | --- |
| Day | Putative Cells | Reads Passing Quality Filter | Reads from putative cells | Mean Reads per cell | Valid Reads aligned to Sample Tags | Reads from putative cells |
| ED18 | 41,998 | 97,33% | 93,87% | 13,258.31 | 5,611,637 | 65,20% |
| d3 | 37,975 | 96,98% | 90,83% | 14,650.78 | 7,944,922 | 68,93% |
| d14 | 54,683 | 96,83% | 92,96% | 12,617.67 | 6,633,932 | 71,24% |

### 3.2 Annotation of lineage defining markers revealed functional states of T cell subclusters

All three timepoints were integrated into a single data set. Clusters of CD3□ T cells were manually annotated based on the expression of lineage-defining marker genes within the integrated dataset. The markers used for cluster identification are listed in Table 3 (see also Supplementary Figure 4). The two largest clusters contained a similar number of cells: n = 13,848 effector αβ T cells (16.96%), defined by the expression of *CD28, ICOS,* and *CD8A*, and n = 13,830 naïve αβ T cells (16.95%), characterized by *FOXO1, KLF2, IL7R, P2PRY8, CD38,* and *RORA*. The largest γδ T cell cluster comprised n = 7,330 mature γδ T cells (8.98%). Annotation of the T cell subclusters revealed distinct functional states across the cell cycle, including differentiating, proliferating, and activated T cells, which collectively contributed to the circular arrangement of clusters in the UMAP representation. Despite prior enrichment for CD3□ T cells, small contaminating populations were consistently detected across most samples, including thrombocytes (n = 1,401 cells; 1.7%), erythrocytes (n = 290 cells; 0.3%), and myeloid cells (n = 138 cells; 0.2%). Thrombocytes were identified by expression of *GP1BB, GP9,* and *F13A1*; erythrocytes by *HBE1, SLC4A1, EPB42, SPTB,* and *DMTN*; and myeloid cells by *ANXA1, CD74, PDCD1LG2, BLB1,* and *BLB2* (Figure 2 A and B; Tables 3–4).

**Figure 2.**
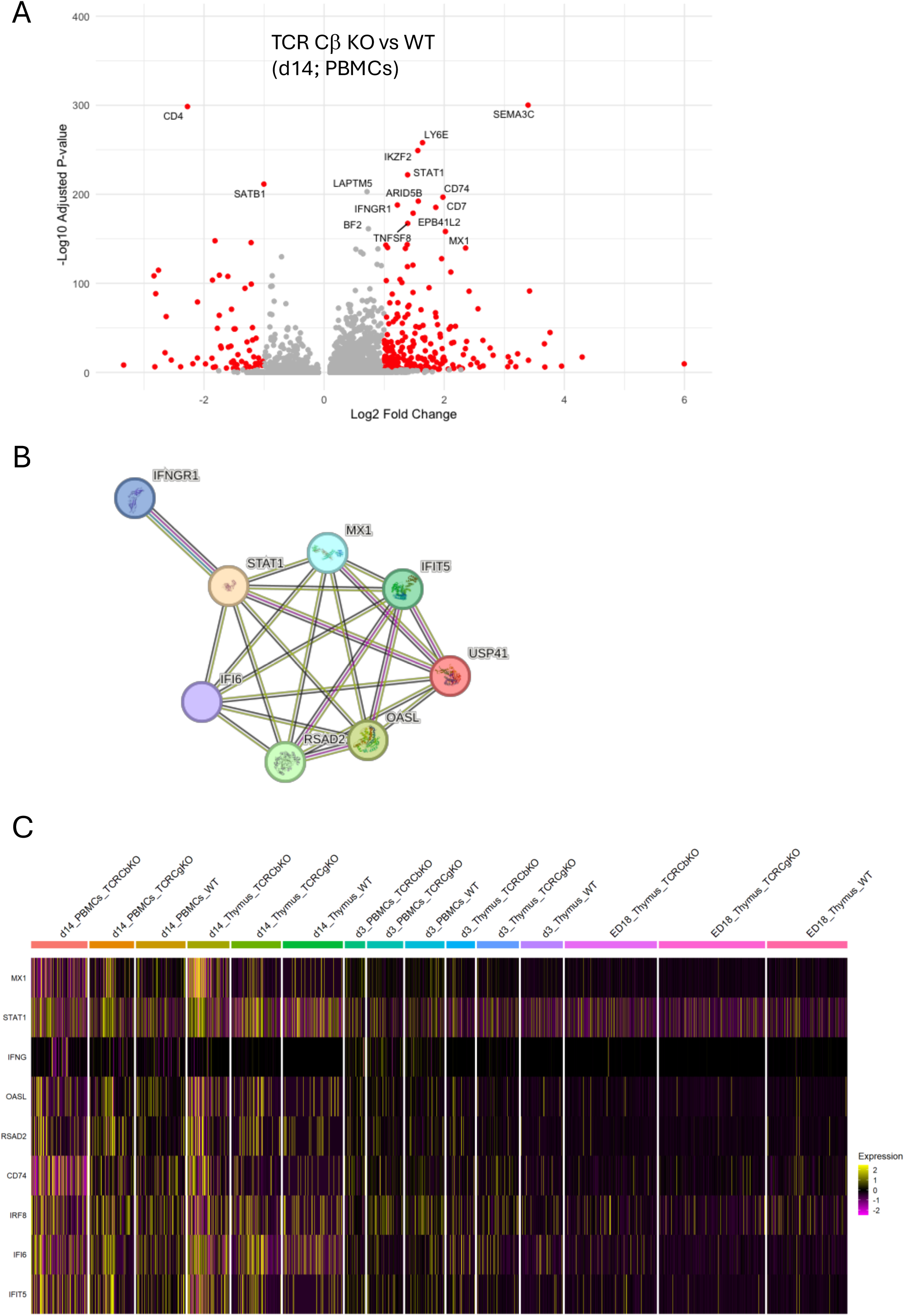
(A) Single-cell RNA sequencing of CD3□T cells isolated from the thymus and PBMCs of WT, TCR Cβ KO and TCR Cγ KO chickens at ED18, d3, and d14 post hatch revealed distinct gene clusters in the integrated UMAP plots. UMAPs showing CD3^+^ T cell clusters annotated according to specific gene markers in Table 4. (B) Percentages (%) of cells in each Cluster (0-17) of integrated data from all three timepoints.

**Table 3.** Cellcount of integrated clusters.

| clusters | Cellcount | Percentage (%) |
| --- | --- | --- |
| 0 | 13848 | 16,96 |
| 1 | 13830 | 16,94 |
| 2 | 10453 | 12,80 |
| 3 | 7330 | 8,98 |
| 4 | 5363 | 6,57 |
| 5 | 5350 | 6,55 |
| 6 | 3539 | 4,34 |
| 7 | 3477 | 4,26 |
| 8 | 3328 | 4,08 |
| 9 | 3139 | 3,85 |
| 10 | 2883 | 3,53 |
| 11 | 2318 | 2,84 |
| 12 | 2149 | 2,63 |
| 13 | 1570 | 1,92 |
| 14 | 1401 | 1,72 |
| 15 | 1229 | 1,51 |
| 16 | 290 | 0,36 |
| 17 | 138 | 0,17 |
| Total | 81636 | 100 |

### 3.3 Subclustering of T cells on ED18 reveals division active precursor T cells

Subclustering the ED18 dataset, TCR Cβ KO chickens exhibited a major subcluster of mature γδ T cells (34,8%, n= 3,313) in the thymus, along with two minor clusters of activated γδ T cells (Cluster 8 =14,4%, n= 1369; Cluster 15= 0,34%, n=32 cells) and a minor cluster of proliferating γδ T cells (10,4%, n = 992 cells). Additionally, division-active precursor (3,4%, n=320 cells) and differentiating (4,7%, n= 447 cells) αβ T cells were observed. In TCR Cγ KO chickens, effector (31,7%, n= 3483) and differentiating (45,4%, n= 4979) αβ T cells constituted major subpopulations together with the division-active precursor cells (8,5%, n= 938 cells). Proliferating αβ (5,1%, n= 561) T cells were also observed. In the wild type siblings, the largest three clusters were also effector T cells (23,5%, n= 1959), differentiating αβ T cells (19,5%, n= 1626) and mature γδ T cells (16,9%, n= 1412), which is comparable to the largest clusters in the TCR Cβ KO and TCR Cγ KO chickens (Figure 3A).

**Figure 3.**
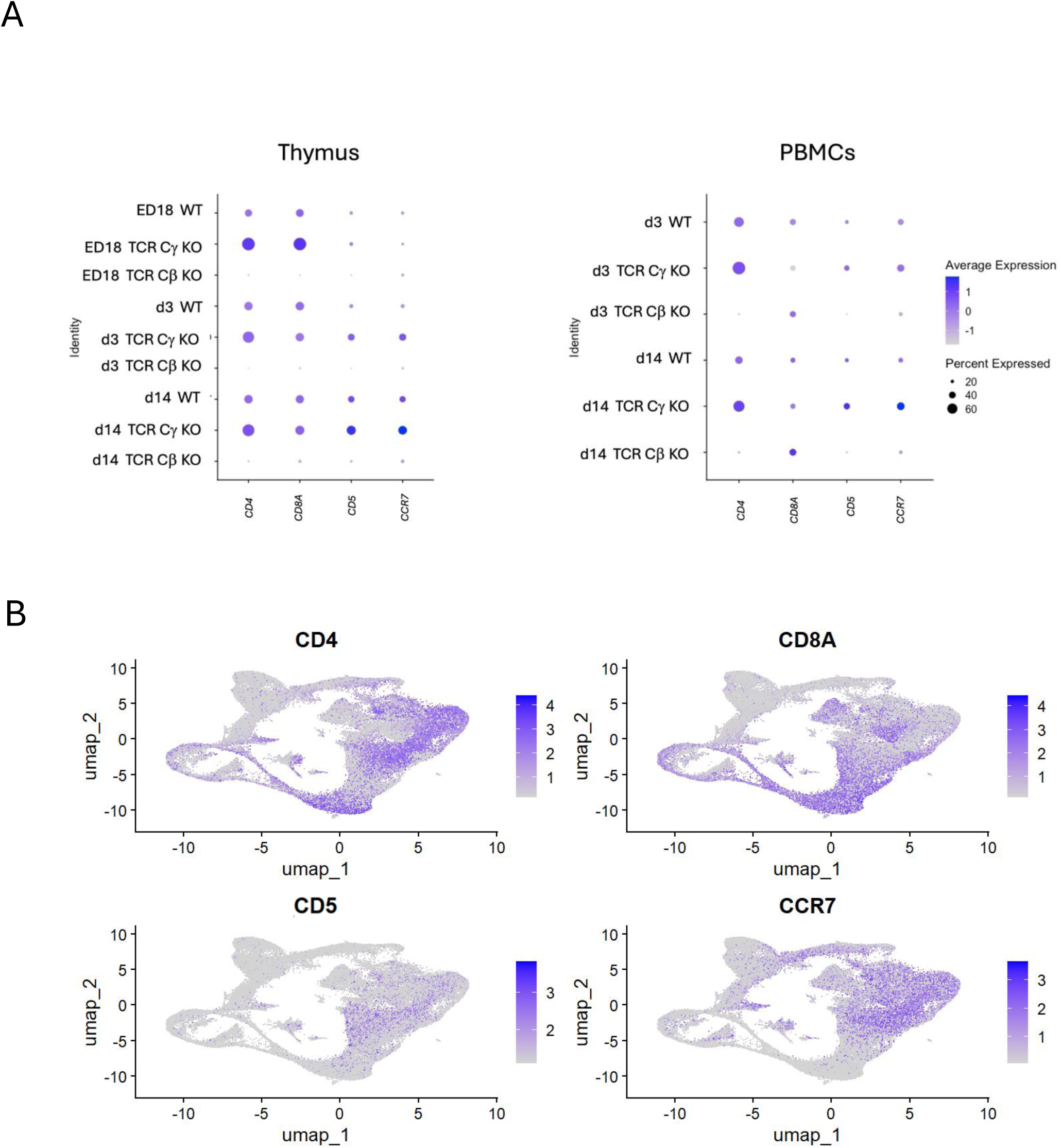
Identification of T cell subsets among chicken CD3^+^ clusters on (A) ED18, (B) d3 and (C) d14 post hatch from WT, TCR Cβ KO and TCR Cγ KO chickens from thymus and PBMCs. UMAPs showing CD3^+^ T cell clusters annotated according to specific gene markers in Table 4.

**Table 4.**
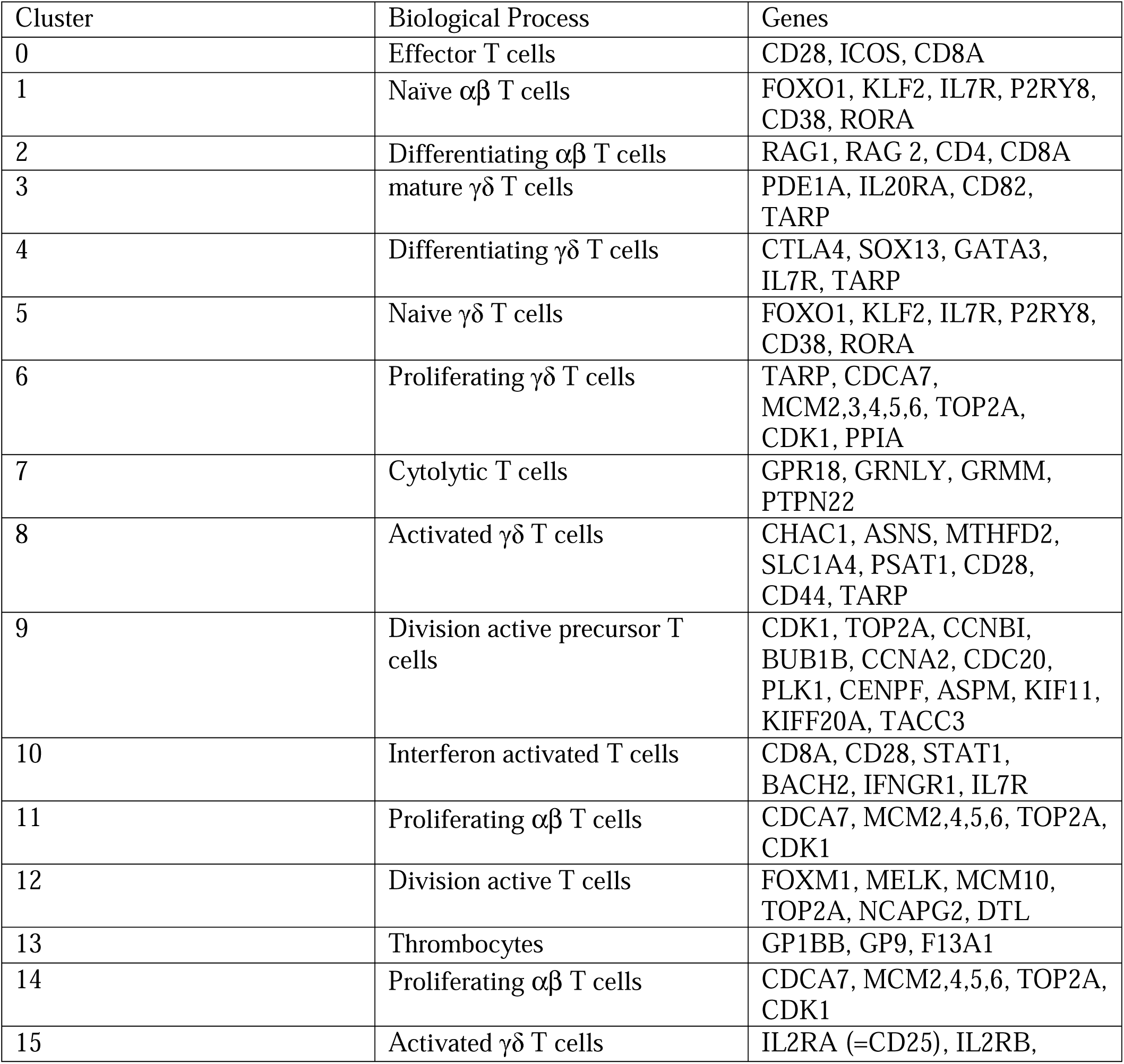
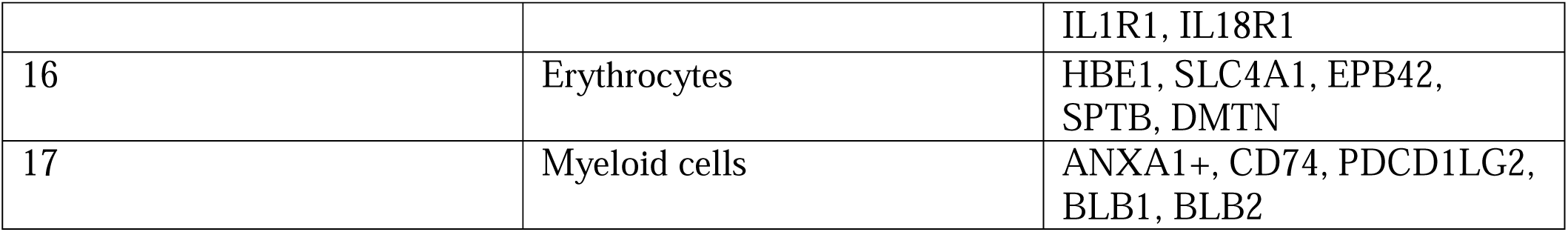
Lineage defining genes for cluster annotation.

### 3.4 Subclustering on d3 shows activated γδ T cells in PBMCs of TCR Cβ KO chickens

Subclustering data from d3 revealed in the thymus of WT chickens, the major clusters contained effector αβ (33,4%, n= 1459) and differentiating αβ T cells (17,0%%, n= 744), naïve T cells (7,8%, n= 341) and mature γδ T cells (8,0%, n= 352). Minor clusters were proliferating αβ (3,6% n= 158) and proliferating γδ (5,7%, n= 249) T cells. In the thymus of TCR Cβ KO the main clusters were also mature γδ T cells (25,1% n=737) and proliferating γδ T cells (12,8%, n=375). But other than in WT also activated γδ T cells (17,3%, n=507) were a dominant population in the thymus of TCR Cβ KO chickens. In the TCR Cγ KO chickens the principal cell type were as in WT chickens effector αβ T cells (40,7%, n=1764), naïve αβ T cells (19,8%, n= 859) and differentiating αβ T cells (9,9%, n=429).

While in PBMCs WT chickens exhibited major clusters of mature αβ T cells (43,8%, n= 1784), naïve T cells (13,0% n= 531) and differentiating γδ T cells (12,9%, n= 525). In PBMCs of TCR Cβ KO birds a principal group of differentiating γδ T cells (PBMCs: 22,9%, n= 482) and naive T cells (26,2%, n= 552), with a minor cluster of cytolytic T cells (6,7%, n=142) were designated. By comparing the PBMCs of TCR Cγ KO and WT chickens, TCR Cγ KO birds displayed an increased population of naïve T cells (57,7%, n= 2159) (Figure 3B).

### 3.5 Cluster annotation reveals different activated γδ T cell clusters in TCR Cβ KO chickens on d14

Next, the data from d14 was subclustered. Comparing TCR Cγ KO and WT T cell clusters in the chicken’s thymus on d14 the clusters of effector αβ T cells (WT: 26,9%, n= 1676; TCR Cγ KO: 30,2%, n= 1544), differentiating αβ T cells (WT: 12,4%, n= 773; TCR Cγ KO: 11,0%, n= 560), and naïve T cells (WT: 12,2%, n= 762; TCR Cγ KO: 22,1%, n= 1130) were mostly similar. In the thymus of TCR Cβ KO chickens mature (18,0%, n=780), differentiating (10,6%, n= 461) and interferon activated γδ T cells (10,2%, n=442) formed major clusters. The PBMCs of TCR Cγ KO chickens showed one dominant cell population: naïve αβ T cells (54,9%, n=2542). Both, TCR Cβ KO and WT chickens, have equal clusters of differentiating γδ T cells (TCR Cβ KO: 23,0%, n= 1175; WT: 19,9%, n= 1018) and naïve T cells (TCR Cβ KO: 19,9%, n= 921; WT: 31,5%, n= 1611) on d14 in the PBMCs. Surprisingly, in TCR Cβ KO chickens we identified two large clusters of activated T cells. The first cluster expressed genes associated with the interferon pathway, including *MX1, STAT1,* and *IFNGR1*, and was therefore designated as interferon-activated γδ T cells (12.0%, n= 520). The second group, which represented the major population in TCR Cβ KO chickens at day 14, consisted of cytolytic T cells characterized by the expression of *GZMM* and *GNLY* (31.0%, n= 1,345) (Figure 3C).

### 3.6 TARP as a useful marker to identify γδ T cell clusters

To describe γδ T cells in more detail, *TARP* was identified for γδ T cells as a potential marker to annotate γδ T cell clusters since it showed high expression (>1) in over 75% of the cells from TCR Cβ KO chickens at every timepoint in both PBMCs and thymus. In wild type *TARP* was intermediately expressed (=0) in around 50% of the cells and had a negative expression in TCR Cγ KO chickens (< -1) at all three timepoints (Figure 4A). *MAF, SOX13, GATA3, KK34, INPP5A, PDE1A, CD82* and *IL20RA* showed equal high expression in TCR Cβ KO chickens and intermediate expression in WT chickens. However, the relative expression of *SOX13, GATA3, PDE1A* and *CD82* were higher compared to *MAF, KK34, INPP3A* and *IL20RA*. Leading to the conclusion that these genes were found in specific γδ T cells clusters only. Feature Plot indicated the location of cells expressing genes describing γδ T cells on the integrated UMAP. While *TARP, GATA3* and *CD82* seem to be expressed in all γδ T cell clusters, *PDE1A, KK34, INPP5A* and *IL20RA* are only expressed in specific γδ T cell clusters (Figure 3B). *MAF* was only expressed in activated and naïve T cells and was not exclusively expressed in γδ T cells but also αβ T cells. *KK34* was only expressed in naïve γδ T cells. Alongside, the markers *PDE1A* and *CD82* were highly expressed in the γδ T cells. However, only *PDE1A* seemed to be completely absent in TCR Cγ KO chickens (Figure 4B). To identify a large population of αβ T cells *CD4, CD8A, CD5 and CCR7* were used. All four genes were absent in TCR Cβ KO chickens in the thymus. In PBMCs we found high expression in around 20% of the cells from TCR Cβ KO chickens at d3 and d14, indicating differentiated CD8^+^ γδ T cells. Moreover, high expression (>1) of *CD4* and *CD8A* in 40-60% of the cells in the thymus of TCR Cγ KO chickens at all timepoints was found and intermediate expression in WT chickens at all timepoints in the thymus (Figure 5A). While *CD4* and *CD8A* are present in both naïve and activated αβ T cells, *CD5* was present in effector αβ T cells and *CCR7* was used to identify naïve αβ T cells. (Figure 5B).

**Figure 4.**
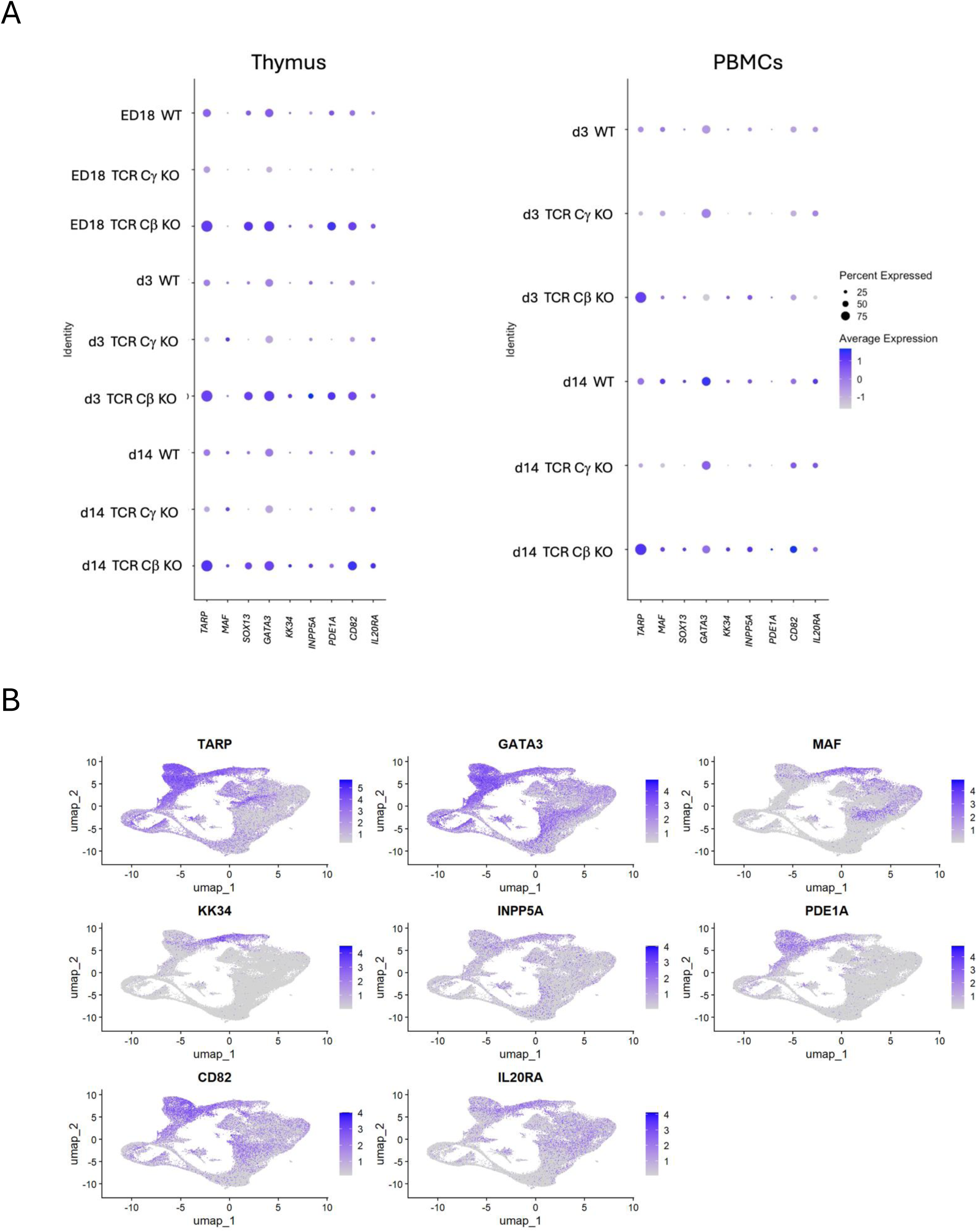
Identification of markers for γδ T cell subsets among chicken CD3^+^ cells. (A) Dot plot representing the expression of a selection of γδ T cell associated genes in subgroups grouped by timepoint, organ and genotype. The radius of the dot corresponds to the percentage of cells in each cluster expressing the gene, and color intensity corresponds to scaled expression values (average_log2 fold change). (B) Feature plot of potential γδ T cell markers in integrated UMAP cluster. The color intensity corresponds to scaled expression values.

**Figure 5.**
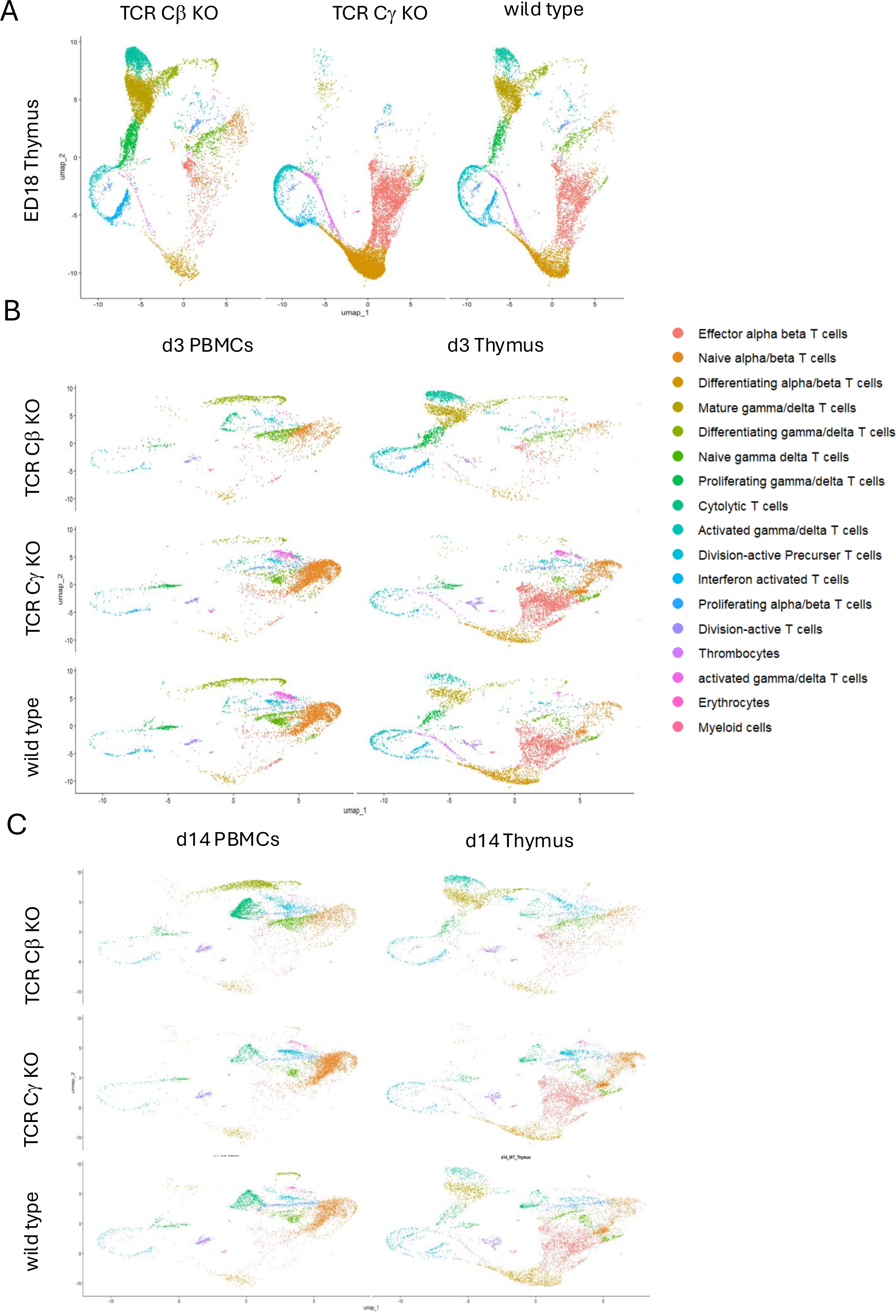
Identification of markers for αβ T cell subsets among chicken CD3^+^ cells from PBMCs. (A) Dot plot representing the expression of a selection of αβ T cell associated genes in in subgroups grouped by timepoint, organ and genotype. The radius of the dot corresponds to the percentage of cells in each cluster expressing the gene, and color intensity corresponds to scaled expression values (average_log2 fold change). (B) Feature plot of potential αβ T cell markers in integrated UMAP cluster. The color represents the expression level (average_log2 fold change).

### 3.7 Interferon pathway activated in TCR Cβ KO chickens

Comparing genes which are significantly upregulated in TCR Cβ KO compared to WT chickens at d14 after hatch, revealed genes which are involved in the interferon pathway (Figure 6A). The string networks showed the close relation of the genes involved into interferon pathway, such as *STAT1, MX1* and *OASL* (Figure 6B). A heatmap analysis of genes that are involved in the interferon pathway (*STAT1, IFI6, MX1, USP41, IFNGR1, OASL, RSAD2, IFIT5)* showed strong expression (>2) of these genes in the thymus and PBMCs of TCR Cβ KO chickens on d14. While there are less cell clusters in WT and TCR Cγ KO chickens on d14 that express genes involved in the interferon pathway.

**Figure 6.**
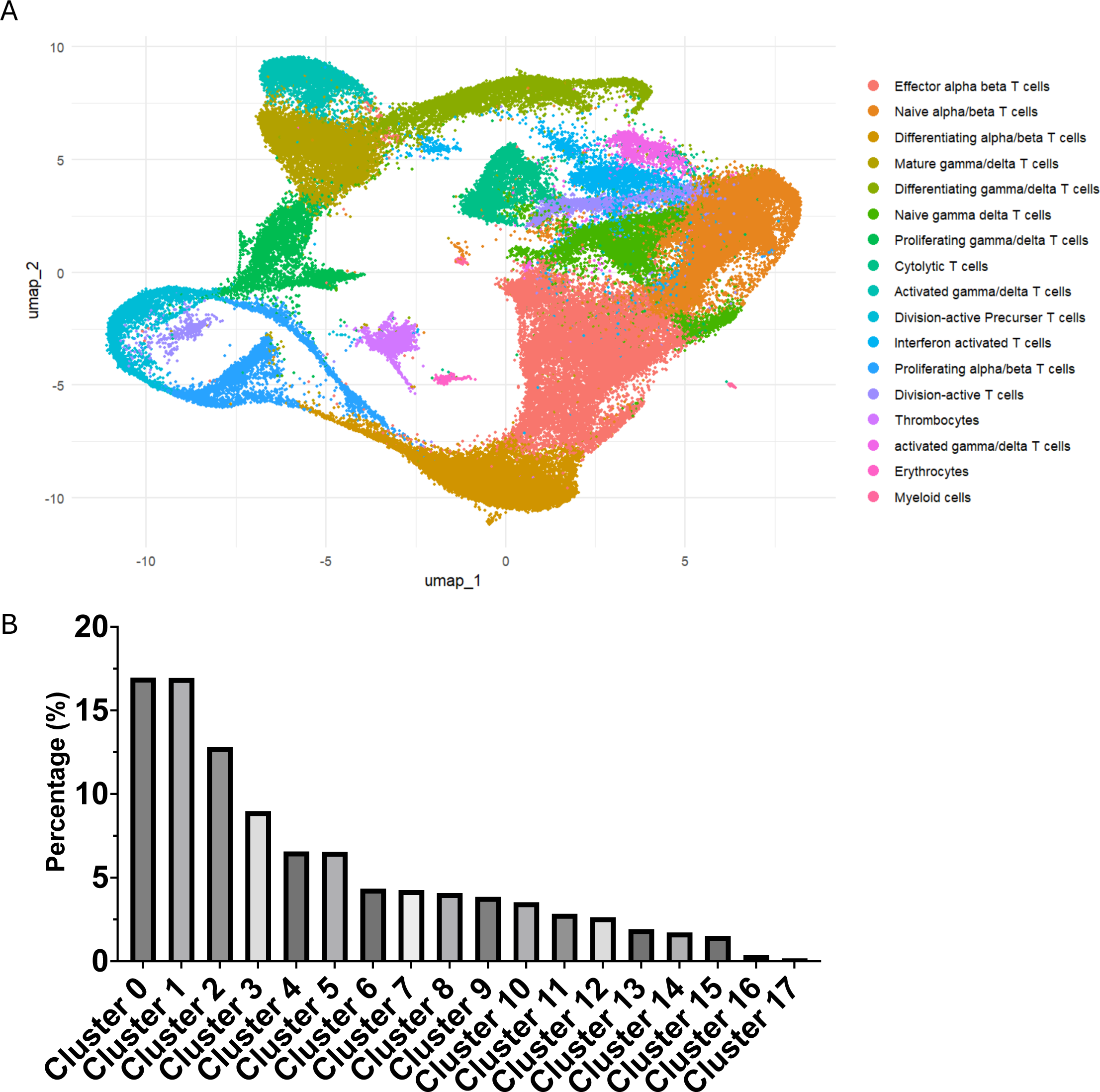
Interferon-activated γδ T cell clusters. A) Volcano plot of differentially expressed genes (DEGs) between TCR Cβ KO and WT chickens in PBMCs at d14 post hatch. The log2(FC) indicates the mean fold-change (FC) in expression levels for each gene, and each dot represents one gene. DEGs identified in pairwise comparisons between TCR Cβ KO and WT chickens are defined as p < 0.05, FC > |2|. DEGs with higher and lower expression are represented by red dots, while genes with no substantial change in expression are indicated as grey dots. B) String Network of DEGs from TCR Cβ KO compared to WT chickens involved in interferon pathway. C) Heatmap of clusters from thymus and PBMCs of WT, TCR Cβ KO and TCR Cγ KO chickens at d14 showing genes involved in the interferon pathway. The color represents the gene expression level (average_log2 fold change).

Interestingly, in the Thymus and PBMCs on d3 and ED18 less expression of interferon-stimulated genes (Figure 6C) was detected.

### 3.8 Cytolytic CD8+ γδ T cells expressing Granulysin and Granzyme A

The analysis of cytolytic genes on d14 in the T cells showed, that TCR Cβ KO chicken express cytokines that are related to lysis and apoptosis. High expression (n= 2) of Granulysin (*GNLY*), Granzyme A (*GZMA*), Granzyme M (*GZMM*) and *IL2RB* in the PBMCs of TCR Cβ KO chickens was detected on d14, while this was not observed in the thymus of these chickens. Also, on ED18 and d3 there is no such upregulation of cytotoxic γδ T cells visible in any genotype (Figure 7A). The string networks show the close relation of the cytokines and chemokines involved in cytotoxicity (Figure 7B).

**Figure 7.**
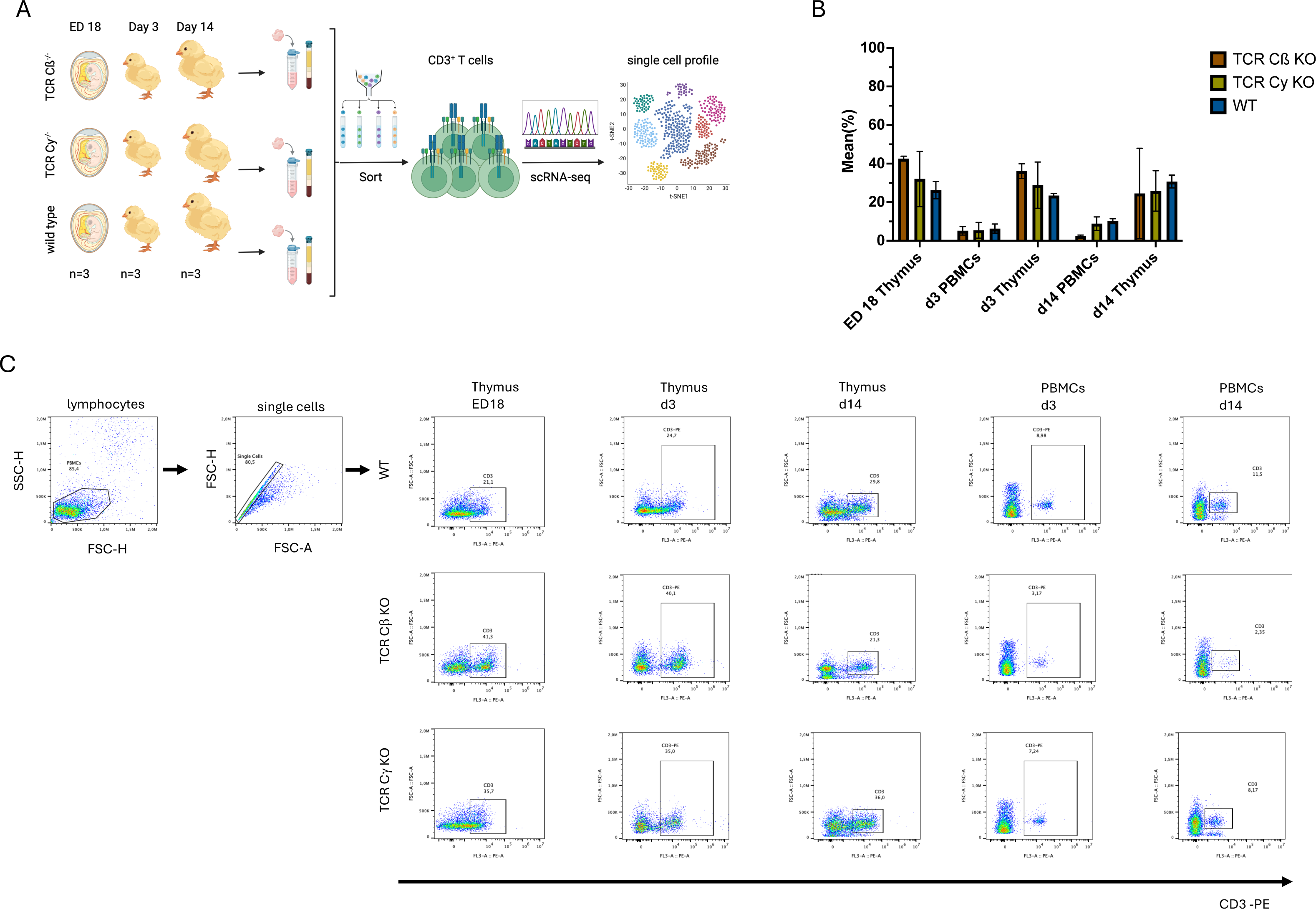
Cytolytic γδ T cell clusters. A) Heatmap of Clusters from thymus and PBMCs of WT, TCR Cβ KO and TCR Cγ KO chickens 14 days after hatch showing genes involved in cytolytic activity. The color represents the gene expression level (average_log2 fold change). B) String Network of DEGs involved in cytolytic activity.

## 4 Discussion

Single cell RNA-sequencing is a novel tool to understand gene expression patterns in cells to define clusters (Tang, Barbacioru et al. 2009, Eschke, Moore et al. 2023). In this study, we used WTA to analyze T cell clusters in TCR KO and WT chickens from embryonic day 18 (ED18) until two weeks post hatch (d14).

Data integration was performed across all genotypes (TCR Cβ KO, TCR Cγ KO, and wild type), developmental stages (ED18, d3, and d14), and organs (thymus and PBMCs). This approach enables a more comprehensive comparison across the different layers of the analysis. However, integrating all datasets also introduces the risk of misclassification of a subset of single cells. Although T cells can be subdivided into αβ and γδ lineages, both share common transcriptional programs related to processes such as proliferation and differentiation. As a result, cells from different T-cell subgroups may cluster together despite belonging to distinct lineages (Hao, Hao et al. 2021)

The first T cells that move into the thymus are TCR1^+^ cells which occur first on ED12 and are first detectable in the spleen by ED15. The second wave of T cells colonize the thymus at ED15 and move towards spleen on ED19 (Chen, Sowder et al. 1989, Coltey, Bucy et al. 1989, Chen, Gobel et al. 1994). Even though there should be some CD3^+^ T cells in the blood of ED18, we were not able to sort enough cells for scRNA-seq.

Through the use of TCR KO chicken, it was possible to detect different genes that are specific markers for γδ or αβ T cells. Before, *TARP, MAF, GATA3, KK34* and *INPP5A* were described as γδ T cell markers (Maxwell, Soderlund et al. 2024). Likewise, in this study we also found that *TARP* and *GATA3* are highly expressed in the γδ T cells. While *MAF* is only expressed in activated and mature T cells and is not exclusively expressed in γδ T cells but also αβ T cells. The genes *KK34* and *INPP5A* show a low expression but are exclusively expressed in γδ T cells. Alongside we found that the markers *PDE1A* and *CD82* are highly expressed in the γδ T cells. However, only *PDE1A* seems to be absent in TCR Cγ KO chickens, while *TARP* and *CD82* is also expressed in some cells of the TCR Cγ KO chickens. *IL20RA* showed low expression but is a selective marker for the detailed description of γδ T cells. *IL20RA* is known to be a subunit of the IL20 receptor and is involved in the regulation of inflammatory mediators by regulating the *JAK-STAT* pathway (Lamichhane, Mo et al. 2021). *CD82* (also known as *KAI1* or *R2*) is an inducible membrane marker that was initially identified from a T cell activation study (Gaugitsch, Hofer et al. 1991). Here, we highlight for the first time, that *CD82* is not exclusively expressed in γδ T cells, but comparing TCR Cβ KO with TCR Cγ KO and WT chickens, we see that most of γδ T cells express *CD82* while only few αβ T cells express *CD82*. Until now there is no literature on γδ T cells expressing the calmodulin-dependent phosphodiesterase 1A (*PDE1A*) exclusively. This study reveals for the first time the absence of *PDE1A* in TCR Cγ KO chickens and shows that only a specific γδ T cell cluster express *PDE1A*.

Before, scRNA-seq in chicken was performed in peripheral blood leukocytes (Maxwell, Soderlund et al. 2024). Here, the T cell clusters were subdivided in CD4, T_regs_, CD8, γδ T cells and cytolytic cells. However, in our study we subdivided the T cells not only by the markers from Maxwell et al (2024) but also further due to their differentiation status into proliferation, differentiation, naïve T cells and mature T cells (Figure 2,3). Likewise, in comparison to Maxwell et al. we found cytolytic T cells, that show high expression of Granulysin and Granzyme A and M in TCR Cβ KO animals 14 days after hatch (Maxwell, Soderlund et al. 2024). Other than shown before, here we can show that this cytolytic T cells are γδ T cells in TCR Cβ KO birds. Interestingly, these cytolytic γδ T cells do neither express perforin nor Granzyme B.

Depending on the expression of specific cytokines, helper T cells are subdivided in Th1, Th2, Th17 and T_regs_ in humans and mammals (Raphael, Nalawade et al. 2015). Still, it was not possible to define these T cell subsets for the helper T cells in chickens due to missing annotations of some cytokines and transcription factors. In this analysis, the T cells were subdivided into different clusters depending on their activation status. However, we did not find individual clusters for different helper T cells subclass, since in this study the sorted CD3^+^ T cells in the wildtype showed no expression of IFN-γ (Th1), IL-4 (Th2), IL-17 (Th17) or IL-10 (T_reg_). Even though these markers are already annotated in the chicken genome.

In addition to T cell populations, we identified minor clusters of thrombocytes (expressing *GP1BB, GP9*, and *F13A1*), erythrocytes (expressing *HBE1, SLC4A1*, and *EPB42*), and myeloid cells (expressing *BLB1* and *BLB2*) (see Table 4; Figures 2 and 3), despite the fact that samples were sorted for CD3 □T cells. These clusters are clearly distinct from the T cell populations, represent less than 2% of all cells, and therefore do not interfere with downstream analyses. Maxwell et al. (2024) used an additional immunomagnetic separation step to reduce thrombocytes based on the expression of CD41/61. However, even using a cleanup step still a population of 5-7% Thrombocyte was measured in their samples (Maxwell, Soderlund et al. 2024). These irregularities do not interfere with the T cell analysis, as these cell clusters can be clearly annotated using specific markers.

In the thymus at both ED18 and d3, we identified two clusters (clusters 8 and 15) that we annotated as activated γδ T cells. These clusters shared core γδ T cell markers such as *TARP, ZAP70*, and *CD28*, supporting their classification. Despite this common identity, they exhibited distinct activation marker profiles. Cluster 8 expressed *CHAC1, ASNS, MTHFD2, SLC1A4, PSAT1,* and *CD44*, while cluster 15 was characterized by the expression of *IL2RA (CD25), IL2RB, IL1R1,* and *IL18R1* (see Tables 4, Figures 2 and 3). Although the activation markers differed between the two clusters, both were classified broadly as activated γδ T cells, as a more refined subclassification was not feasible based on the available data.

In this study, we identified for the first time a subset of γδ T cells that exhibit the expression of genes associated with the interferon pathway, including *MX1*, *STAT1*, and *OASL* (Mountford, Gheyas et al. 2022). TCR Cβ KO chickens display a severe phenotype as early as 14 days post hatch, characterized by inflammation of the gut, spleen, and skin. This condition is driven by an imbalance between humoral and cytotoxic responses within the adaptive immune system (von Heyl, Klinger et al. 2023). The interferon-activated γδ T cells described here might play a defining role in the severity of the phenotype observed in TCR Cβ KO chickens. Further studies are needed to determine whether these T cells are also present in other inflammatory diseases in chickens and to assess if their differentiation is triggered by the activation of dysregulated monocytes.

## 5 Conclusion

This study revealed that cytolytic T cells, characterized by the expression of Granulysin and Granzymes, are predominantly γδ T cells in TCR Cβ KO chickens. Both the cytolytic γδ T cells and the interferon-activated γδ T cells are likely contributors to the severe phenotype observed in TCR Cβ KO chickens. However, the precise mechanisms underlying this phenotype remain unclear, highlighting the need for future sequencing analyses of monocytes and B cells to further elucidate the drivers of this response.

Overall, this study provides new insights into chicken T cell biology, including T cell markers for specific subsets such as αβ T cells and γδ T cells, as well as their functional states, such as proliferation and differentiation. Notably, the use of TCR Cβ KO chickens has uncovered novel characteristics of γδ T cells, offering valuable perspectives for further research.

## 6 List of abbreviations

αβ: alpha beta
γδ: gamma delta
B cell: B Lymphocyte
CD: Cluster of differentiation
D: day
DEG: Differentially expressed gene
ED: Embryonal day
FBS: Fetal bovine serum
G: Gauge
GO: Gene ontology
KO: Knockout
PBMC: Peripheral blood mononuclear cells
PBS: Phosphate-buffered saline
RPE: random priming and extension
T cell: T Lymphocyte
TCR: T cell receptor
TCR Cβ KO: Chicken line lacking alpha beta T cells
TCR Cγ KO: Chicken line lacking gamma delta T cells
scRNA-seq: Single cell RNA-sequencing
UMAP: Uniform Manifold Approximation and Projection
WT: Wild type
WTA: Whole transcriptomic analysis
°C: Degrees Celsius

## 7 Ethics approval and consent to participate

Animal experiments were approved by the government of Upper Bavaria, Germany (ROB-55.2-2532.Vet_02-17-101 & 55.2-1-54-2532-104-2015). Experiments were performed according to the German Welfare Act and European Union Normative for Care and Use of Experimental Animals.

## 8 Consent for publication

Not applicable

## 9 Data Availability Statement

Raw data for this study can be found on SRA under the accession number PRJNA1226579.

## 12 Author Contributions

TvH performed and analyzed the experiments and wrote the paper. BS supervised the work and wrote the paper. MB and SPF helped with the analysis of the data. AS and CW performed experiments. All authors approved the submitted version.

## 13 Acknowledgments

ChatGPT (https://chat.openai.com/) was used to improve conciseness of language and coding, with no contribution to the scientific content.

