## Supplementary Information for "Unraveling new characteristics of γδ T cells using scRNA-seq in TCR KO chicken"

### *Supplementary Material*

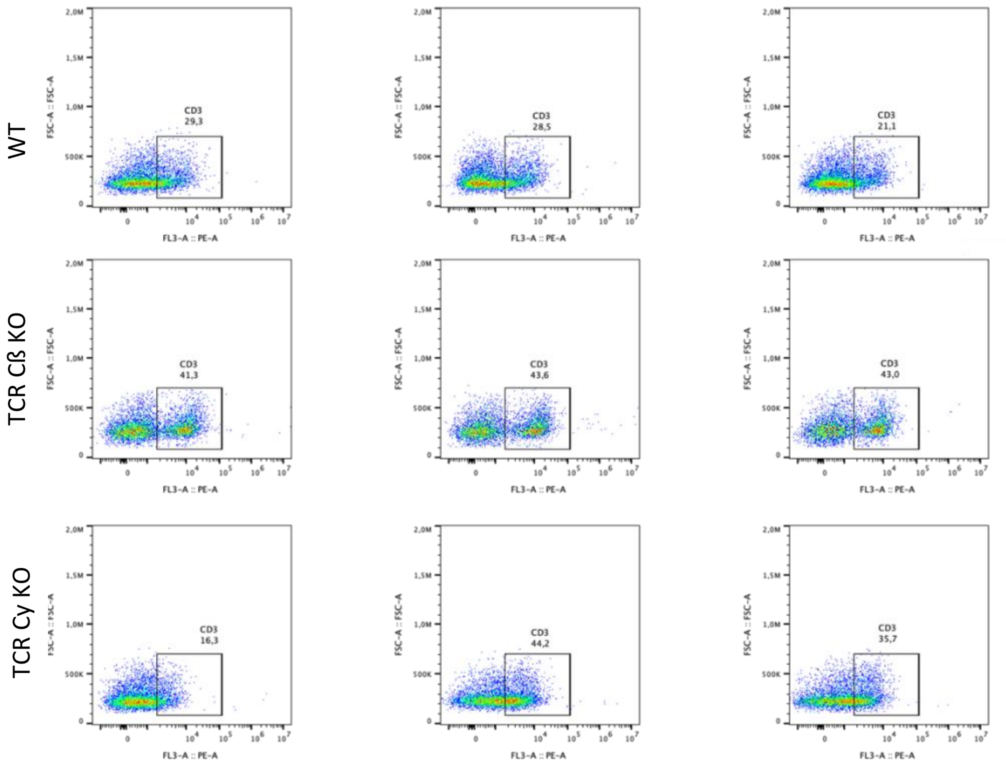


**Supplementary Figure 1.** Schematic representation of the flow-cytometry gating strategy for $CD3^+$ cell percentages in thymus. Flow-cytometry analysis of $CD3^+$ (CD3-PE) within single cells in percentages from wild type, TCR Cβ KO and TCR C γ KO at ED18.

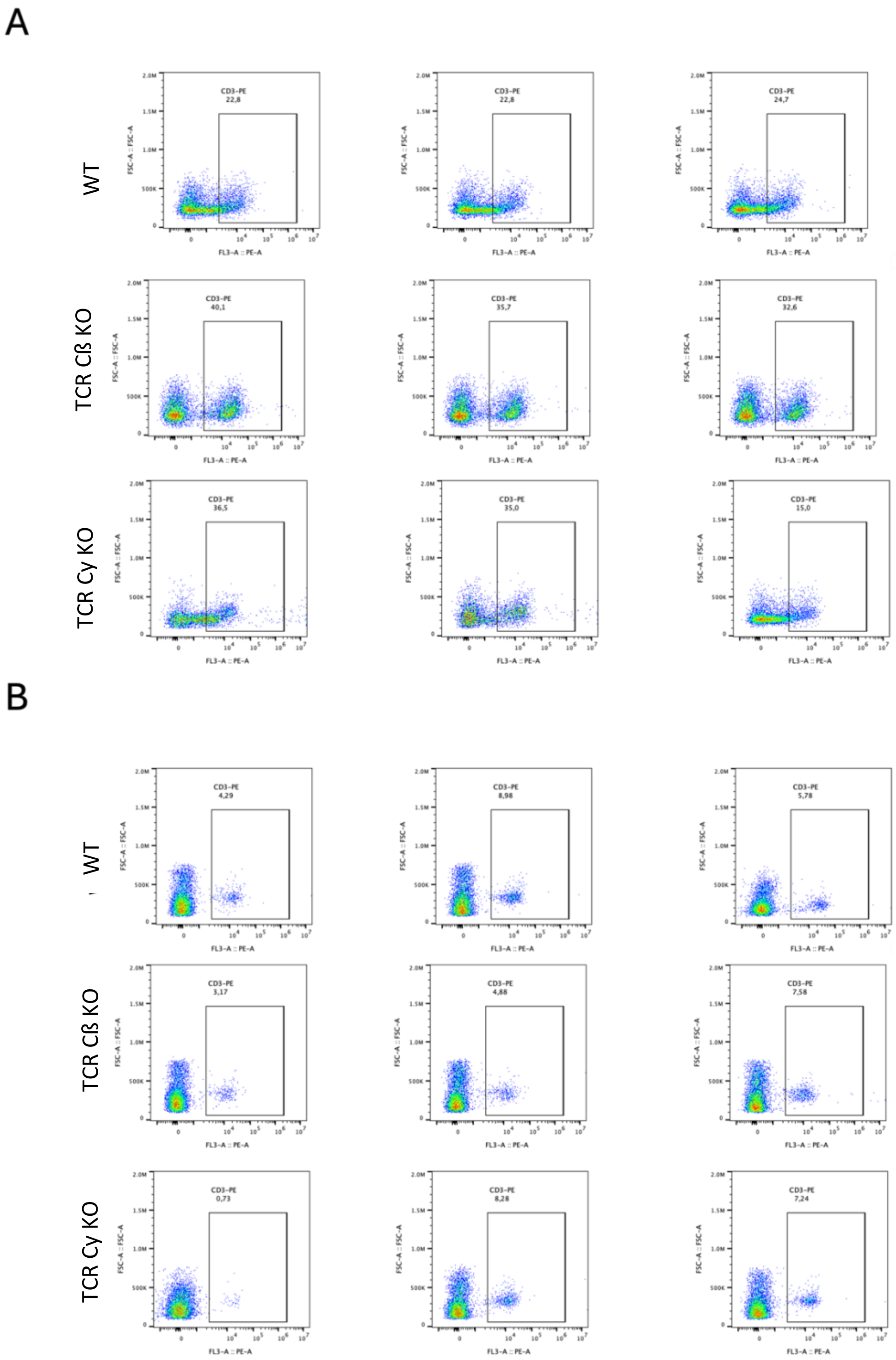


**Supplementary Figure 2.** Schematic representation of the flow-cytometry gating strategy for $CD3^+$ cell percentages in A) thymus and B) PBMCs. Flow-cytometry analysis of $CD3^+$ (CD3-PE) within single cells in percentages from wild type, TCR Cβ KO and TCR C γ KO at d3.

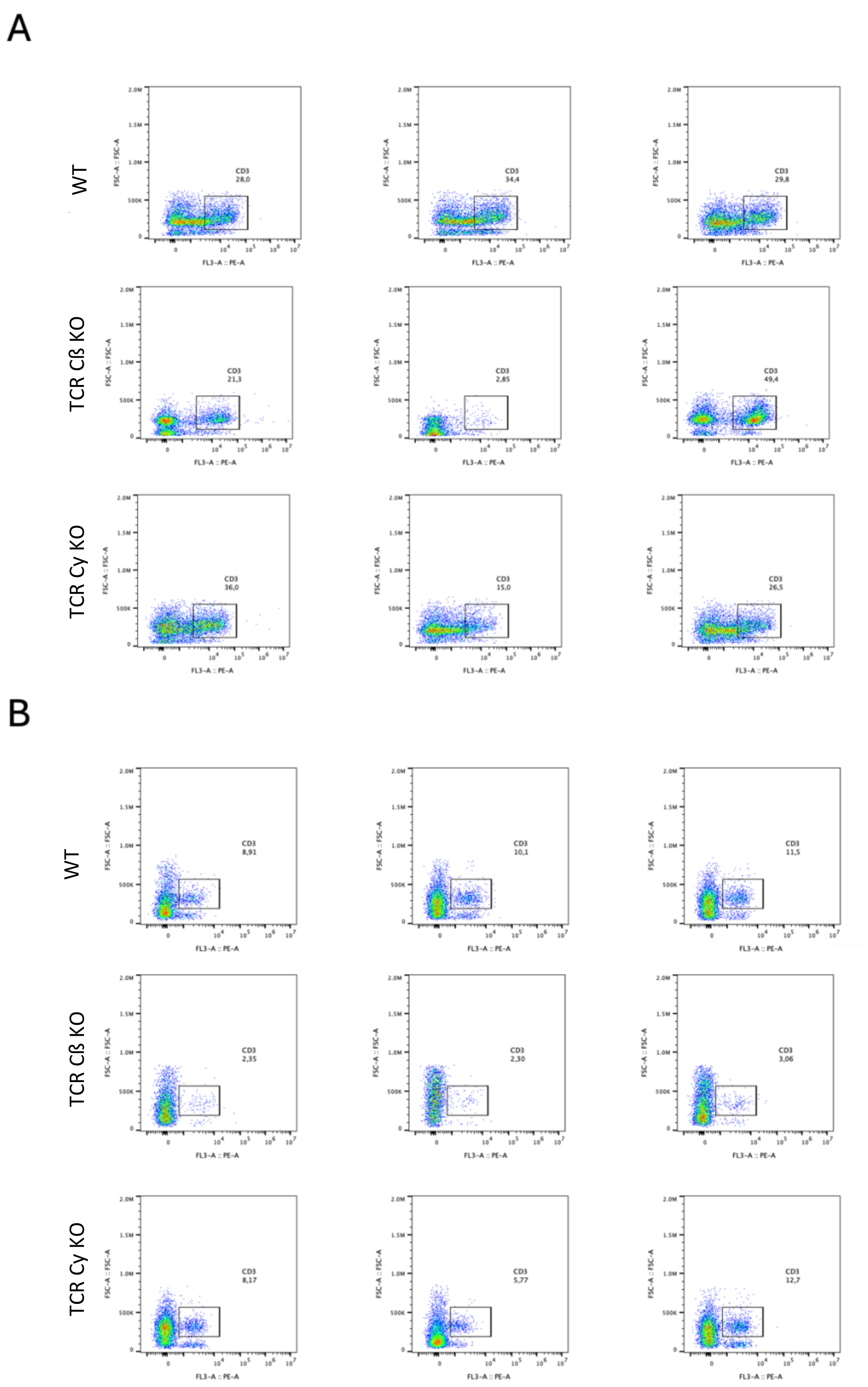


**Supplementary Figure 3.** Schematic representation of the flow-cytometry gating strategy for CD3$^+$ cell percentages in A) thymus and B) PBMCs. Flow-cytometry analysis of CD3$^+$ (CD3-PE) within single cells in percentages from wild type, TCR Cβ KO and TCR C γ KO at d14.

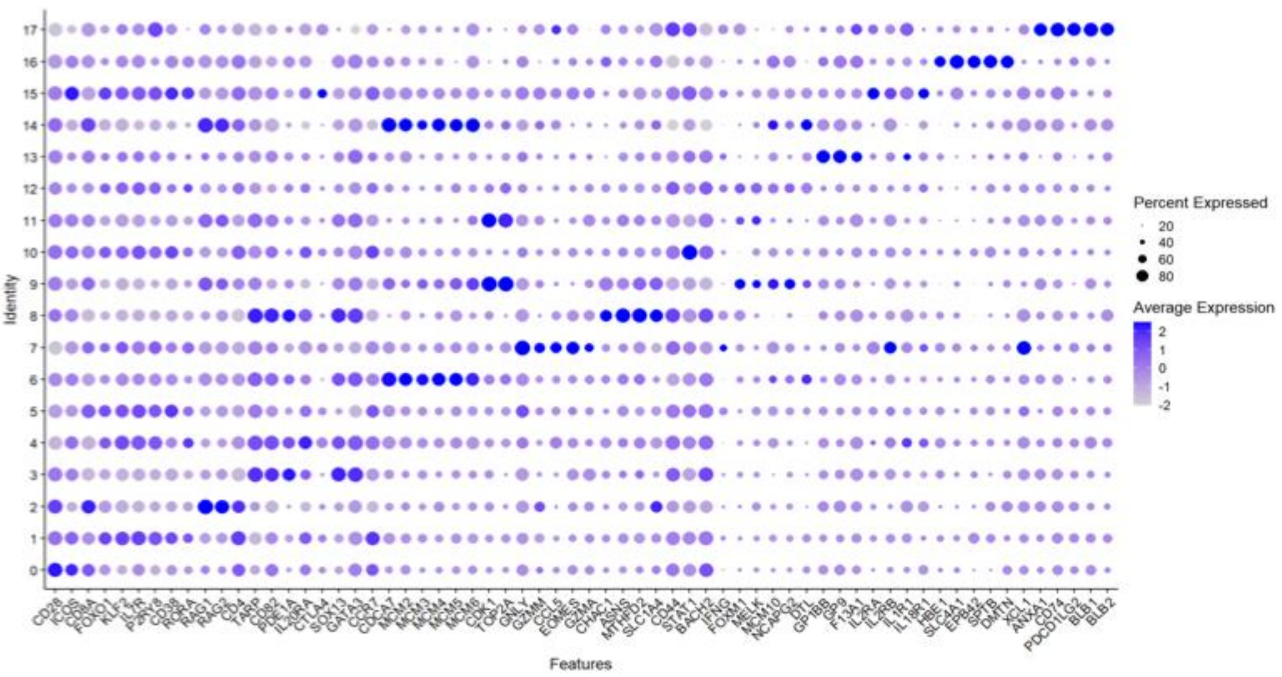


**Supplementary Figure 4.** Dot plot representing the expression of a selection of T cell associated genes in the integrated Clusters 0-17. The radius of the dot corresponds to the percentage of cells in each cluster expressing the gene, and color intensity corresponds to scaled expression values (average_log2 fold change).

**Supplementary Table 1. Cellcount and percentages of each cluster from TCR Cβ KO, TCR Cγ Ko and WT animals on ED18, d3 and d14 in thymus**

| Cluster | Cellcount | Group |
|---|---|---|
| 0 | 131 | d14_TCRCbKO_PBMCs |
| 1 | 921 | d14_TCRCbKO_PBMCs |
| 2 | 140 | d14_TCRCbKO_PBMCs |
| 3 | 35 | d14_TCRCbKO_PBMCs |
| 4 | 1175 | d14_TCRCbKO_PBMCs |
| 5 | 819 | d14_TCRCbKO_PBMCs |
| 6 | 135 | d14_TCRCbKO_PBMCs |
| 7 | 1345 | d14_TCRCbKO_PBMCs |
| 8 | 7 | d14_TCRCbKO_PBMCs |
| 9 | 37 | d14_TCRCbKO_PBMCs |
| 10 | 520 | d14_TCRCbKO_PBMCs |
| 11 | 64 | d14_TCRCbKO_PBMCs |
| 12 | 177 | d14_TCRCbKO_PBMCs |
| 13 | 178 | d14_TCRCbKO_PBMCs |
| 14 | 10 | d14_TCRCbKO_PBMCs |
| 15 | 68 | d14_TCRCbKO_PBMCs |
| 16 | 25 | d14_TCRCbKO_PBMCs |
| 17 | 19 | d14_TCRCbKO_PBMCs |
| 0 | 137 | d14_TCRCgKO_PBMCs |
| 1 | 2542 | d14_TCRCgKO_PBMCs |
| 2 | 154 | d14_TCRCgKO_PBMCs |
| 3 | 12 | d14_TCRCgKO_PBMCs |
| 4 | 90 | d14_TCRCgKO_PBMCs |
| 5 | 305 | d14_TCRCgKO_PBMCs |
| 6 | 98 | d14_TCRCgKO_PBMCs |
| 7 | 376 | d14_TCRCgKO_PBMCs |
| 8 | 9 | d14_TCRCgKO_PBMCs |
| 9 | 45 | d14_TCRCgKO_PBMCs |
| 10 | 364 | d14_TCRCgKO_PBMCs |
| 11 | 40 | d14_TCRCgKO_PBMCs |
| 12 | 173 | d14_TCRCgKO_PBMCs |
| 13 | 151 | d14_TCRCgKO_PBMCs |
| 14 | 9 | d14_TCRCgKO_PBMCs |
| 15 | 100 | d14_TCRCgKO_PBMCs |
| 16 | 14 | d14_TCRCgKO_PBMCs |
| 17 | 14 | d14_TCRCgKO_PBMCs |
| 0 | 160 | d14_WT_PBMCs |
| 1 | 1611 | d14_WT_PBMCs |
| 2 | 155 | d14_WT_PBMCs |

| | | |
|---|---|---|
| 3 | 38 | d14_WT_PBMCs |
| 4 | 1018 | d14_WT_PBMCs |
| 5 | 381 | d14_WT_PBMCs |
| 6 | 138 | d14_WT_PBMCs |
| 7 | 661 | d14_WT_PBMCs |
| 8 | 5 | d14_WT_PBMCs |
| 9 | 83 | d14_WT_PBMCs |
| 10 | 208 | d14_WT_PBMCs |
| 11 | 50 | d14_WT_PBMCs |
| 12 | 294 | d14_WT_PBMCs |
| 13 | 184 | d14_WT_PBMCs |
| 14 | 11 | d14_WT_PBMCs |
| 15 | 87 | d14_WT_PBMCs |
| 16 | 25 | d14_WT_PBMCs |
| 17 | 11 | d14_WT_PBMCs |
| 0 | 76 | d3_TCRCbKO_PBMCs |
| 1 | 552 | d3_TCRCbKO_PBMCs |
| 2 | 57 | d3_TCRCbKO_PBMCs |
| 3 | 14 | d3_TCRCbKO_PBMCs |
| 4 | 482 | d3_TCRCbKO_PBMCs |
| 5 | 415 | d3_TCRCbKO_PBMCs |
| 6 | 47 | d3_TCRCbKO_PBMCs |
| 7 | 142 | d3_TCRCbKO_PBMCs |
| 8 | 2 | d3_TCRCbKO_PBMCs |
| 9 | 23 | d3_TCRCbKO_PBMCs |
| 10 | 86 | d3_TCRCbKO_PBMCs |
| 11 | 38 | d3_TCRCbKO_PBMCs |
| 12 | 31 | d3_TCRCbKO_PBMCs |
| 13 | 73 | d3_TCRCbKO_PBMCs |
| 14 | 1 | d3_TCRCbKO_PBMCs |
| 15 | 24 | d3_TCRCbKO_PBMCs |
| 16 | 30 | d3_TCRCbKO_PBMCs |
| 17 | 13 | d3_TCRCbKO_PBMCs |
| 0 | 145 | d3_TCRCgKO_PBMCs |
| 1 | 2150 | d3_TCRCgKO_PBMCs |
| 2 | 52 | d3_TCRCgKO_PBMCs |
| 3 | 2 | d3_TCRCgKO_PBMCs |
| 4 | 125 | d3_TCRCgKO_PBMCs |
| 5 | 292 | d3_TCRCgKO_PBMCs |
| 6 | 163 | d3_TCRCgKO_PBMCs |
| 7 | 83 | d3_TCRCgKO_PBMCs |
| 8 | 1 | d3_TCRCgKO_PBMCs |
| 9 | 38 | d3_TCRCgKO_PBMCs |

| | | |
|---|---|---|
| 10 | 113 | d3_TCRCgKO_PBMCs |
| 11 | 63 | d3_TCRCgKO_PBMCs |
| 12 | 126 | d3_TCRCgKO_PBMCs |
| 13 | 79 | d3_TCRCgKO_PBMCs |
| 14 | 1 | d3_TCRCgKO_PBMCs |
| 15 | 247 | d3_TCRCgKO_PBMCs |
| 16 | 31 | d3_TCRCgKO_PBMCs |
| 17 | 14 | d3_TCRCgKO_PBMCs |
| 0 | 145 | d3_WT_PBMCs |
| 1 | 1784 | d3_WT_PBMCs |
| 2 | 93 | d3_WT_PBMCs |
| 3 | 11 | d3_WT_PBMCs |
| 4 | 525 | d3_WT_PBMCs |
| 5 | 531 | d3_WT_PBMCs |
| 6 | 173 | d3_WT_PBMCs |
| 7 | 77 | d3_WT_PBMCs |
| 8 | 4 | d3_WT_PBMCs |
| 9 | 39 | d3_WT_PBMCs |
| 10 | 153 | d3_WT_PBMCs |
| 11 | 88 | d3_WT_PBMCs |
| 12 | 68 | d3_WT_PBMCs |
| 13 | 98 | d3_WT_PBMCs |
| 14 | 11 | d3_WT_PBMCs |
| 15 | 252 | d3_WT_PBMCs |
| 16 | 17 | d3_WT_PBMCs |
| 17 | 5 | d3_WT_PBMCs |
| 0 | 154 | d3_TCRCbKO_Thymus |
| 1 | 125 | d3_TCRCbKO_Thymus |
| 2 | 82 | d3_TCRCbKO_Thymus |
| 3 | 737 | d3_TCRCbKO_Thymus |
| 4 | 167 | d3_TCRCbKO_Thymus |
| 5 | 165 | d3_TCRCbKO_Thymus |
| 6 | 375 | d3_TCRCbKO_Thymus |
| 7 | 35 | d3_TCRCbKO_Thymus |
| 8 | 507 | d3_TCRCbKO_Thymus |
| 9 | 203 | d3_TCRCbKO_Thymus |
| 10 | 48 | d3_TCRCbKO_Thymus |
| 11 | 158 | d3_TCRCbKO_Thymus |
| 12 | 72 | d3_TCRCbKO_Thymus |
| 13 | 52 | d3_TCRCbKO_Thymus |
| 14 | 23 | d3_TCRCbKO_Thymus |
| 15 | 23 | d3_TCRCbKO_Thymus |
| 16 | 8 | d3_TCRCbKO_Thymus |

| | | |
|---|---|---|
| 17 | 5 | d3_TCRCbKO_Thymus |
| 0 | 1764 | d3_TCRCgKO_Thymus |
| 1 | 859 | d3_TCRCgKO_Thymus |
| 2 | 429 | d3_TCRCgKO_Thymus |
| 3 | 10 | d3_TCRCgKO_Thymus |
| 4 | 31 | d3_TCRCgKO_Thymus |
| 5 | 211 | d3_TCRCgKO_Thymus |
| 6 | 134 | d3_TCRCgKO_Thymus |
| 7 | 36 | d3_TCRCgKO_Thymus |
| 8 | 10 | d3_TCRCgKO_Thymus |
| 9 | 118 | d3_TCRCgKO_Thymus |
| 10 | 101 | d3_TCRCgKO_Thymus |
| 11 | 62 | d3_TCRCgKO_Thymus |
| 12 | 133 | d3_TCRCgKO_Thymus |
| 13 | 151 | d3_TCRCgKO_Thymus |
| 14 | 55 | d3_TCRCgKO_Thymus |
| 15 | 174 | d3_TCRCgKO_Thymus |
| 16 | 35 | d3_TCRCgKO_Thymus |
| 17 | 22 | d3_TCRCgKO_Thymus |
| 0 | 1459 | d3_WT_Thymus |
| 1 | 341 | d3_WT_Thymus |
| 2 | 744 | d3_WT_Thymus |
| 3 | 352 | d3_WT_Thymus |
| 4 | 68 | d3_WT_Thymus |
| 5 | 198 | d3_WT_Thymus |
| 6 | 222 | d3_WT_Thymus |
| 7 | 14 | d3_WT_Thymus |
| 8 | 168 | d3_WT_Thymus |
| 9 | 249 | d3_WT_Thymus |
| 10 | 49 | d3_WT_Thymus |
| 11 | 158 | d3_WT_Thymus |
| 12 | 64 | d3_WT_Thymus |
| 13 | 97 | d3_WT_Thymus |
| 14 | 132 | d3_WT_Thymus |
| 15 | 37 | d3_WT_Thymus |
| 16 | 16 | d3_WT_Thymus |
| 17 | 4 | d3_WT_Thymus |
| 0 | 606 | ED18_TCRCbKO_Thymus |
| 1 | 382 | ED18_TCRCbKO_Thymus |
| 2 | 439 | ED18_TCRCbKO_Thymus |
| 3 | 3313 | ED18_TCRCbKO_Thymus |
| 4 | 639 | ED18_TCRCbKO_Thymus |
| 5 | 447 | ED18_TCRCbKO_Thymus |

| | | |
|---|---|---|
| 6 | 992 | ED18_TCRCbKO_Thymus |
| 7 | 47 | ED18_TCRCbKO_Thymus |
| 8 | 1369 | ED18_TCRCbKO_Thymus |
| 9 | 320 | ED18_TCRCbKO_Thymus |
| 10 | 74 | ED18_TCRCbKO_Thymus |
| 11 | 535 | ED18_TCRCbKO_Thymus |
| 12 | 216 | ED18_TCRCbKO_Thymus |
| 13 | 3 | ED18_TCRCbKO_Thymus |
| 14 | 104 | ED18_TCRCbKO_Thymus |
| 15 | 32 | ED18_TCRCbKO_Thymus |
| 16 | 2 | ED18_TCRCbKO_Thymus |
| 17 | 1 | ED18_TCRCbKO_Thymus |
| 0 | 3483 | ED18_TCRCgKO_Thymus |
| 1 | 178 | ED18_TCRCgKO_Thymus |
| 2 | 4979 | ED18_TCRCgKO_Thymus |
| 3 | 93 | ED18_TCRCgKO_Thymus |
| 4 | 3 | ED18_TCRCgKO_Thymus |
| 5 | 92 | ED18_TCRCgKO_Thymus |
| 6 | 63 | ED18_TCRCgKO_Thymus |
| 7 | 1 | ED18_TCRCgKO_Thymus |
| 8 | 42 | ED18_TCRCgKO_Thymus |
| 9 | 938 | ED18_TCRCgKO_Thymus |
| 10 | 45 | ED18_TCRCgKO_Thymus |
| 11 | 367 | ED18_TCRCgKO_Thymus |
| 12 | 106 | ED18_TCRCgKO_Thymus |
| 13 | 9 | ED18_TCRCgKO_Thymus |
| 14 | 561 | ED18_TCRCgKO_Thymus |
| 16 | 17 | ED18_TCRCgKO_Thymus |
| 17 | 1 | ED18_TCRCgKO_Thymus |
| 0 | 1959 | ED18_WT_Thymus |
| 1 | 171 | ED18_WT_Thymus |
| 2 | 1626 | ED18_WT_Thymus |
| 3 | 1412 | ED18_WT_Thymus |
| 4 | 300 | ED18_WT_Thymus |
| 5 | 268 | ED18_WT_Thymus |
| 6 | 466 | ED18_WT_Thymus |
| 7 | 18 | ED18_WT_Thymus |
| 8 | 558 | ED18_WT_Thymus |
| 9 | 624 | ED18_WT_Thymus |
| 10 | 86 | ED18_WT_Thymus |
| 11 | 367 | ED18_WT_Thymus |
| 12 | 144 | ED18_WT_Thymus |
| 13 | 5 | ED18_WT_Thymus |

| | | |
|---|---|---|
| 14 | 320 | ED18_WT_Thymus |
| 15 | 17 | ED18_WT_Thymus |
| 16 | 3 | ED18_WT_Thymus |
| 17 | 1 | ED18_WT_Thymus |
| 0 | 409 | d14_TCRCbKO_Thymus |
| 1 | 322 | d14_TCRCbKO_Thymus |
| 2 | 170 | d14_TCRCbKO_Thymus |
| 3 | 780 | d14_TCRCbKO_Thymus |
| 4 | 461 | d14_TCRCbKO_Thymus |
| 5 | 400 | d14_TCRCbKO_Thymus |
| 6 | 187 | d14_TCRCbKO_Thymus |
| 7 | 156 | d14_TCRCbKO_Thymus |
| 8 | 434 | d14_TCRCbKO_Thymus |
| 9 | 104 | d14_TCRCbKO_Thymus |
| 10 | 442 | d14_TCRCbKO_Thymus |
| 11 | 127 | d14_TCRCbKO_Thymus |
| 12 | 95 | d14_TCRCbKO_Thymus |
| 13 | 139 | d14_TCRCbKO_Thymus |
| 14 | 32 | d14_TCRCbKO_Thymus |
| 15 | 47 | d14_TCRCbKO_Thymus |
| 16 | 20 | d14_TCRCbKO_Thymus |
| 17 | 9 | d14_TCRCbKO_Thymus |
| 0 | 1544 | d14_TCRCgKO_Thymus |
| 1 | 1130 | d14_TCRCgKO_Thymus |
| 2 | 560 | d14_TCRCgKO_Thymus |
| 3 | 18 | d14_TCRCgKO_Thymus |
| 4 | 51 | d14_TCRCgKO_Thymus |
| 5 | 331 | d14_TCRCgKO_Thymus |
| 6 | 109 | d14_TCRCgKO_Thymus |
| 7 | 233 | d14_TCRCgKO_Thymus |
| 8 | 7 | d14_TCRCgKO_Thymus |
| 9 | 136 | d14_TCRCgKO_Thymus |
| 10 | 385 | d14_TCRCgKO_Thymus |
| 11 | 74 | d14_TCRCgKO_Thymus |
| 12 | 194 | d14_TCRCgKO_Thymus |
| 13 | 161 | d14_TCRCgKO_Thymus |
| 14 | 66 | d14_TCRCgKO_Thymus |
| 15 | 79 | d14_TCRCgKO_Thymus |
| 16 | 26 | d14_TCRCgKO_Thymus |
| 17 | 11 | d14_TCRCgKO_Thymus |
| 0 | 1676 | d14_WT_Thymus |
| 1 | 762 | d14_WT_Thymus |
| 2 | 773 | d14_WT_Thymus |

| | | |
|---|---|---|
| 3 | 503 | d14_WT_Thymus |
| 4 | 228 | d14_WT_Thymus |
| 5 | 495 | d14_WT_Thymus |
| 6 | 237 | d14_WT_Thymus |
| 7 | 253 | d14_WT_Thymus |
| 8 | 206 | d14_WT_Thymus |
| 9 | 182 | d14_WT_Thymus |
| 10 | 209 | d14_WT_Thymus |
| 11 | 127 | d14_WT_Thymus |
| 12 | 256 | d14_WT_Thymus |
| 13 | 190 | d14_WT_Thymus |
| 14 | 65 | d14_WT_Thymus |
| 15 | 42 | d14_WT_Thymus |
| 16 | 21 | d14_WT_Thymus |
| 17 | 8 | d14_WT_Thymus |
